# A molecular phylogeny of Schizothoracinae (Teleostei: Cypriniformes: Cyprinidae) based on 12 protein-coding mitochondrial genes and RAG1 gene analysis

**DOI:** 10.1101/523639

**Authors:** Yurong Du, Ting Wang, Delin Qi, Desheng Qi, Weilin Li, Jiangbin Zhong, Juan Chen, Songchang Guo, Jianbin Ma

## Abstract

The ever-increasing interest in the investigation of origin and speciation of schizothoracine fishes can be dated to 20th century. However, molecular phylogeny of Schizothoracinae and their phylogenetic relationships, as well as the divergence times still remain controversial. In this study, two DNA sets consisting of 12 protein-coding mitochondrial genes from 254 individuals and RAG1 gene from 106 individuals were used to reconstruct the phylogenetic relationships and calculate the divergence times among the subfamily schizothoracinae. Our results indicated that both of the data sets supported a non-monophyletic relationship due to involving of species of *Barbinae*. However, the phylogenetic relationships based on mtDNA genes were more reliable than that inferred from RAG1 gene. The highly specialized grade formed a monophyletic group, together with *Ptychobarbus* as a sister group of *Diptychus* and *Gymnodiptychus*, which was belonging to specialized grade, indicating that *Ptychobarbus* may be transition species to involve to highly specialized schizothoracianae. In addition, the primitive grade clustered with *Percocypris pingi*, a species of *Barbinae*. Based on mtDNA gene, the speciation time of Schizothoracinae was 66 Ma, and the divergence time of the primitive grade and *Percocypris pingi* was 64 Ma. The speciation times of the three grades Schizothoracinae were 57 Ma, 51 Ma and 43 Ma, respectively; and the divergence time of specialized and highly specialized grade was 46 Ma. The divergence times of three grades were not consistent with the three stages of uplift of Qinghai-Tibet Plateau, which is older than the times.

---

Qinghai-Tibet Plateau, with an average altitude of 4000 meters, is the largest, youngest and highest Plateau in the word (Zhang Ti-Cao et al., 2013), which is called the roof of the world (Cao et al., 1981). The environment and climate of Qinghai-Tibet Plateau had changed drastically due to the intensive uplift of the Plateau since Quaternary. Its extreme environment (hypothermia and hypoxia) has been pregnant with rich and colorful life, making it to be one of the world’s biodiversity hotspots. The uplift of Qinghai-Tibet Plateau had experienced three stages in the Eocene, the Oligocene-Miocene, and the Pliocene-Quaternary, respectively (Paul et al., 2001). The first stage of uplift, making Tibet Plateau reached 3000m elevation, happened in late Eocene when India began to wedge into Eurasia (Dewey et al., 1989). From 30Ma to 10Ma, the second phase of uplift occurred. However, at this stage, the Qinghai-Tibet Plateau proceeded with pediplanation cycle (an equilibrium of gradual uplifting, faulting and erosion) as a plain (Dewey et al., 1989). In the late Miocene, pediplanation cycle was ended with faulting and uplifting of the Tibetan Plateau excessive erosion, and at the beginning of Pliocene, the Plateau resumed uplift until Quaternary. At present, there is no accurate time or process of the Qinghai-Tibet Plateau uplift because of the complexity of the process.

Schizothoracine fishes, commonly known as “mountain carps” and characterized by low growth rate, low fecundity and late sexual maturity (Chen Z.M. & Chen, 2001), belong to Cyprinidae, Cypriniformes, Teleostei and are endemic, typical to the Qinghai-Tibet Plateau and its circumjacent areas (Cao et al., 1981). The name of Schizothoracinae came from their characters of the two rows of large and ordered anal scales, locating on both sides of the anus and anal fin that made the blank between the two rows of anal scales like a crack (Wu Y.F. & Wu, 1992). This subfamily includes 15 genera and more than 100 species all over the world (Mirza, 1991), and most of Schizothoracine fishes, 11genera and at least 76 species and subspecies distribute in the whole of China (Chen Y.F. & Cao, 2000; Wu Yun-Fei & Tan, 1991). As their major habitat, nine genera and more than half of species and subspecies distribute in the main drainages of Qinghai-Tibet Plateau of China such as Yellow River, Yangtze River, Yarlung Zangbo River, Irrawaddy River and Qinghai Lake (Cao et al., 1981).

As the extreme environment of Qinghai-Tibet Plateau, the mitochondrial genome is well suited for analysis the phylogenetic relationships for that energy metabolism, depending on the mitochondria, is the most apparent response to the adaption to hypothermia and hypoxia. And then, length of the sequence is necessary for maximum likelihood analysis to clarify interrelationships amongst long-diverged groups on account of that ML trees converge to the true tree as the number of sites approaches infinity (Rogers, 2001), and the general length of mitochondrial genomes is 1.6kb, which is sufficient for the phylogenetic tree. The problem of limitation in the number of sequence sites can be settled by the entire mitochondrial genomes (Inoue J.G. et al., 2003; Inoue Jun G. et al., 2004; Ioue et al., 2001; Ishiguro et al., 2003; Kawahara et al., 2008; Lavoue et al., 2008; Lavoue et al., 2005; Masaki et al., 2004; Minegishi et al., 2005; Miya Masaki & Nishida, 1999; Miya M. & Nishida, 2000; Miya Masaki et al., 2003; Saitoh et al., 2003; Saitoh et al., 2000; Saitoh et al., 2006). However, using mtDNA alone as the molecular marker can only reveal a single linkage group with one history while the utilization of multiple nuclear sequences often shows a conflict among gene trees due to hybridization, lineage sorting, paralogy or selection (Graham et al., 2017). Hence, both the mitochondrial genome and the nuclear gene were used in this study to avoid the possible differences.

Their special living environment makes them form the characters which are highly adapted to hypothermia, hypoxia and high radiation. As for those characters, Schizothoracine fishes have become a good model of Qinghai-Tibet Plateau to study the mechanism adapting to the extreme environment. Based on the morphological characters, Cao (Cao et al., 1981) divided the schizothoracinae into three grades named morphological primitive schizothoracine fishes, morphological specialized schizothoracine fishes, and morphological highly specialized schizothoracine fishes. The primitive fishes, including *Racina, Schizothorax* and *Aspiorhynchus*, are covered by fine scales and these fishes have three rows of pharyngeal teeth and two pairs of barbell. Most of the specialized fishes’ scales tend to degenerate and have two rows of pharyngeal teeth and one pair of barbell, and *Ptychobarbus, Diptychus*, and *Gymnodiptychus* are included in specialized grade. The highly specialized fishes’ scales completely disappear and have only one row of pharyngeal teeth and have no barbell; highly specialized group consists of six genera named *Gymnocypris, Oxygymnocypris*, *Schizopygopsis, Platypharodon, Chuanchia*, and *Herzenstein*. The statistical studies showed a relationship between species richness and elevation that the primitive groups peaked in the low elevation areas of 1250m to 2500m; the specialized fishes mainly distributed in the elevation of 2750m to 3750m and highly specialized grade fishes primary occupied the elevation zone of 3750m to 4750m (Cao et al., 1981). Some researches revealed that the speciation of three grades schizothoracine fishes resulted from the three stages of uplift of Qinghai-Tibet Plateau, which caused the branch of river basins.

By now, there are still many controversies on the phylogeny of Schizothoracinae and their close species. A large body of researches showing that Schizothoracinae originate from the primitive genera of Barbinae, such as *Barbodes, Varicorhinus, Barbus*, based on molecule or morphology (Cao et al., 1981; Chen Y.F. & Cao, 2000; Wu Xianwen et al., 1981), is widely accepted. Gaubert et al. (Gaubert et al., 2009) repute that Schizothoracinae’s closest relatives to be the Barbinae fishes, which is consistent with He (He et al., 2004) and Qi (Qi et al., 2013; Qi et al., 2012), and the subfamily Schizothoracinae is not a monophyly with supertree from the Cyprinidae family level (Gaubert et al., 2009), which is the same to Li (Li Y.L. et al., 2013), but contradict the results of He (He et al., 2004) and Qi (Qi et al., 2012). In the study of Ruber in 2007 indicate that Barbine sensu stricto is closely related to Schizothoracinae, but it’s an unresolved trifurcation between Barbine sensu stricto and two lineages of Schizothoracinae (Ruber et al., 2007). In the research of Li, however, Barbus barbus is embedded in Schizothoracinae that between the specialized group and the primitive group as the sister clade of specialized fishes with the method of mitochondrial genomes of three species of Schizothoracinae and single cytb gene (Li Y.L. et al., 2013). It’s clear that Percocypris pingi, which belongs to *Percocypris* genus and Barbinae subfamily, cluster with the primitive clade in Wang’s study (Wang et al., 2013). Many reasons could explain the inconsistency between morphological phylogeny and molecular phylogeny. For example, He considers the morphological characters may don’t fellow the evolutionary trace of the groups but are shaped by adapting to their survival environmental condition (He & Chen, 2007), such as convergent evolution (Qi et al., 2012). In addition, the low level of sequence divergence and ancestral polymorphism will make it difficult to discriminate species (He & Chen, 2007). The number of samples and the length or quantity of genes also play an important role in the reconstruction of phylogeny. For above reasons, using the molecular method, sufficient samples and appropriate gene to analysis the phylogenetic relationships of Schizothoracinae is becoming more and more urgent.

The overwhelming majority of experts agree with that the speciation of three grades Schizothoracinae fishes results from the uplift of Qinghai-Tibet Plateau that causing the branch of those watersheds. At the middle Tertiary-Dingqing group(late Miocene or early Pliocene), with the movement of the Himalayas, Barbinae, being adapted to the warm climate, gradually tend to adapted to cold weather, which lead to speciate the Plesioschizothorax macrocephalus, which is the only fossil of Schizothoracinae in the world (Wu Y.F. & Wu, 1992). Based on geology and paleontology, Wu deduced that in the first phase of the Himalaya movement, as the early stage of Schizohtoracinae, Plesioschizothorax macrocephalus occurred. In the second phase of the Himalaya movement, the environment was so suitable for freshwater that Plesioschizothorax macrocephalus became the main fishes of those lakes, making this phase to be boom period. However, in the third phase of Himalaya movement, in which the crust began to lift and lakes began to shrink and separate, radical changes of environment resulted in the Plesioschizothorax macrocephalus’ differentiation, migration and extinction. Except Li used the partial mitochondrial genome of *Cytb* gene to declare that the primitive clade divided in the late Eocene and the Rapid speciation events of each clade from the Late Miocene to the Pliocene, corresponding to the time of the geologic acceleration of the Qinghai-Tibetan Plateau (Li Y.L. et al., 2013), there is no researches about the divergence time of Schizothoracine fishes from the subfamily level. Hence it’s essential to deduce the divergence time of three grades of Schizothoracinae on the subfamily level to make up the blank of this area.

In order to better resolve the problems of phylogenetic relationships of Schizothoracinae and calculate the divergence times of these genera, we used 12 protein-coding mitochondrial genes from 254 individuals, which were belonging to 115 species, 11 genera, and RAG1 genes from 106 individuals, which were belonging to 42 species, 8 genera, to reconstruct the phylogenetic tree of the subfamily Schizothoracinae, and inferred the divergence times among Schizothoracine fishes, and analyzed the relationships between the differentiation of Schzithoracinae and the uplift of Qinghai-Tibet Plateau.

## Materials and methods

### Taxon sampling and DNA extraction

We employed 254 mtDNA sequences (table 1) belonging to 11genera of subfamily Cyprinidae, 32 sequences among which were previous work of our team (GenBank accession number: KT833082-KT833113). 4 sequences of Ictiobus were selected as outgroups. The other mitochondrial genomes were downloaded in NCBI (http://www.ncbi.nlm.nih.gov/) and the Genbank accession numbers were shown in Supplementary Material S1.

For the RAG1 gene, 38 individuals were collected by ourselves from the rivers of Qinghai Province and their tributary. We took the muscles or fins and then immediately preserved in 95% ethanol stored at −20 °C for DNA extraction. Total genomic DNA was isolated by using phenol-chloroform extraction (Sambrook et al., 1989) and adjusted to 100ng/mL after testing its concentration on a NanoDrop 2000 supermicro spectrophotometer (Thermo Fisher Scientific Inc., USA). Meanwhile, 68 sequences of RAG1 gene were obtained from GenBank (Accession numbers and species name were showed in Supplementary Material S2).

### PCR Amplification and Sequencing

The RAG1 gene was amplified using the pair of primers RF (5’-CTG AGC TGC AGT CAG TAC CAT AAG ATG T-3’) and RR (5’-TGA GCC TCC ATG AAC TTC TGA AGR TAY TT-3’) (Saitoh & Chen, 2008). The PCR (polymerase chain reaction) reacted at 50uL total volume as follows: 15uL H_2_O, 25uL 2 × PCR Master Mix buffer (Novoprotein Science and Technology Ltd., Shanghai, China), 2.5uL each primer (10mM), and 5uL genomic DNA. And the PCR reaction performed at initial denaturation step at 94 °C for 4 min followed by 35 cycles at 94 °C for 40s, 59°C for 40s, and 72 °C for 90s; with a final extension at 72 °C for 10min. 2μl of amplified DNA was fractionated by electrophoresis through 1% agrose gels. And those PCR products with clear single band were sent to sequence in both directions in Beijing Biomed Company (Beijing, China).

### Sequence editing and analysis

The RAG1 gene sequences were assembled using Contigexpression v9.1.0 software and aligned by online MAFFT version 7 (http://mafft.cbrc.jp/alignment/server/), and then carefully checked by eye and edited manually in MAGE 6.0.

We used 12 protein-coding regions of mtDNA for sequence analysis, which the genes were encoded on the heavy-strand. Then the indices of substitution saturation for these sequences were estimated using DAMBE 5 software with the GTR model (http://dambe.bio.uottawa.ca/software.asp).

### Phylogenetic analysis

There were 254 sequences of mitochondrial genomes using in phylogenetic analysis and 4 common sucker mitochondrial genomes were designed as outgroups (Saitoh et al., 2011b) (Supplementary Material S1). Meanwhile, a total of 106 sequences of RAG1 gene was used and 1 common sucker RAG1 sequence was designed as the outgroup (Supplementary Material S2).

We used 12 protein coding regions to phylogenetic analysis, which the genes were encoded on the heavy-strand. The ND6 wasn’t included in the analysis because this gene was encoded on the light-strand that was different from the 12 genes. All of these sequences were aligned by online MAFFT version 7, and corrected by eyes. A few parts were ambiguous that were excluded. We also removed start codons, stop codons and the overlapping regions between the coding genes ATP6-ATP8, ATP6-COXIII, ND4L-ND4, and ND5-ND6 (Li Y.L. et al., 2013).

The model selection was implemented in Modeltest v3.7. GTR+I+G, as the best model of mtDNA sequences, was used to construct the phylogenetic trees, while SYM+I+G was selected as the best model of RAG1 sequences. GTR+I+G was used as the best model of RAG1 for subsequent analysis because there was no SYM+I+G model in the analysis software. We reconstructed the phylogenetic trees with maximum likelihood (ML) method and Bayesian inference (BI) method. The ML analysis was accomplished by RAxML v7.2.8 (Stamatakis et al., 2008), with GTRGAMMA and GTRCAT models. The BI method was used MrBayes v3.2.4 (Huelsenbeck & Ronquist, 2005; Ronquist & Huelsenbeck, 2003). We used the GTR model to run 2,000,000 generations of 4 simultaneous Monte Carlo Markov chains (MCMC). The sampling frequency and pint frequency were set 100, the diagnostic frequency was set 1000, and the genes were divided into 12 partitions and the likelihood scores lower than those at saturation (burn-in = 500) trees were discarded from the analysis. The posterior probabilities (BBP) of nodes were estimated based on the 50% majority rule consensus of the trees.

### Divergence time estimation

Based on mtDNA genes, we used Beast v2.3.0 (Bouckaert et al., 2014) to calculate the divergence times of three grades of Schizothoracinae with the lognormal incorrect relaxed clock and GTR+G+I model. We used “Speciation: Yule Process” to run 500,000,000 chains. The calibrated nodes and constraint were as followed: basal to *Cyprinus* (33.9Ma) (Saitoh et al., 2011a), and *Labeo bata* / *Labeo senegalensis* (49.1 Ma - 75.1 Ma) (Li Y.L. et al., 2013). And then convergence diagnostics were examined with Tracer v1.6 (http://tree.bio.ed.ac.uk/software/tracer/).

## Results

### Mitochondrial genomes features

A total of 254 mtDNA and 106 RAG1 sequences were obtained after manually editing, which were 10791bp and 1314bp in length, respectively. The average base composition of 12 protein-coding genes was A=28.0%, T=27.8%, G=16.4% and C=27.8%. The base composition of schizothoracinae showed the same characters with teleost fishes that A+T content is significantly higher than the G+C content with an obvious anti-G bias (Jiang et al., 2009; Jondeung et al., 2007; Tzeng et al., 1992). Additionally, the average base composition of RAG1 gene was A=25.3%, T=24.3%, G=26.5% and C=23.9%. There was no obvious base bias among RAG1 gene sequences.

Saturation analyses showed that neither of the mtDNA sequences nor RAG1 sequences was saturated and variable sites of RAG1 sequences were obviously distributed at codon 3 (Figure 1.). Thus, all sequences could be used in reconstructing phylogenetic trees for schizothoricine fishes.

**Figure 1.**
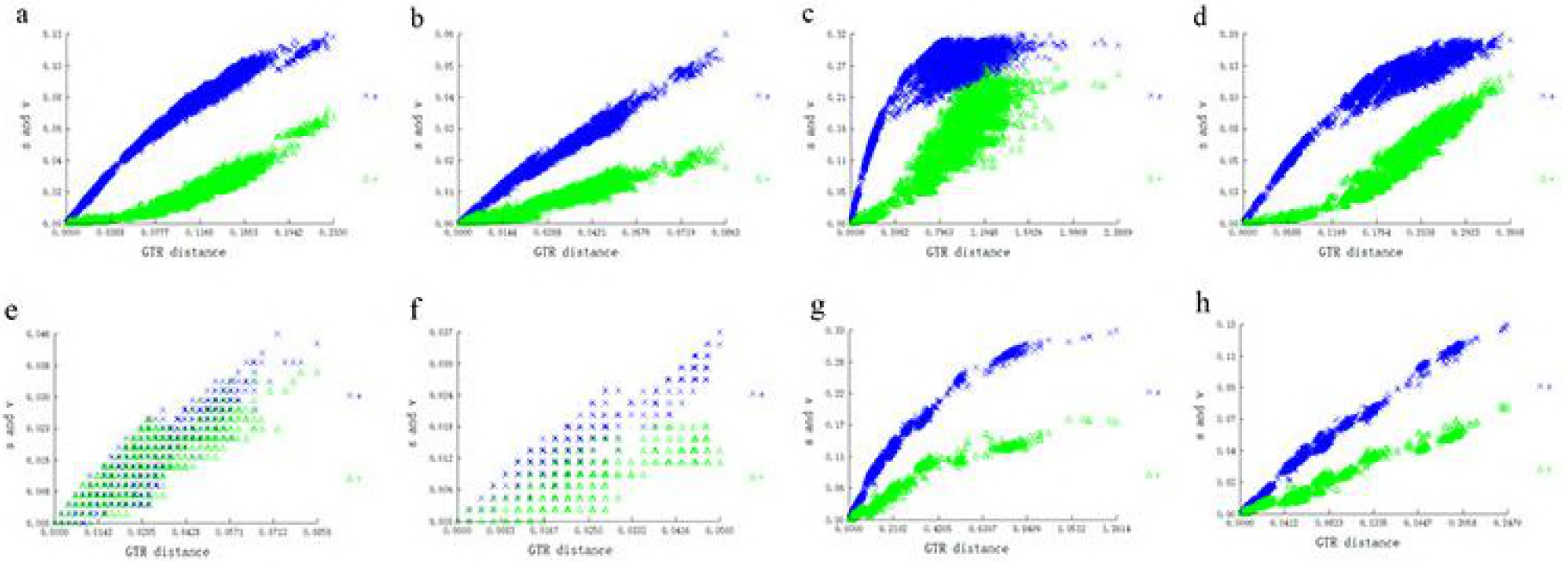
The saturation analysis of both the mitochondrial protein-coding genes (excluding ND6 and 12SrRNA) sequences and RAG1 gene sequences based on the GTR model. Figure a, b, c, and d represent saturation of codon1, codon2 codon3 and complete codon of 12 protein-coding genes, respectively; Figure e, f, g, and h represent saturation of the same part of RAG1 gene.

### Phylogenetic analysis

The maximum likelihood trees inferred from mitochondrial genomes of Cyprinidae were almost the same, only ML tree based on GTRCAT model was shown in Fig 2 (C), which was consistent with the Bayes tree Fig 2 (A). Similarly, the ML trees of RAG1 gene were almost the same, ML tree based on GTRCAT model showed in Fig 2 (D) was also consistent with BI tree(Fig 2 B). The trees based on RAG1 gene indicated that the Schizothoracinae was not a monophyletic groups, which the primitive grade clustered into a big branch with node support ratio of 98 and clade that include *Barbus barbus* and *Scaphiodonichthys acanthopterus* clustered with specialized and highly specialized clade with node support ratio of 37, indicating that the specialized and highly specialized schizothoracinae, as well as the species of Barbinae, as well as the species of *Barbinae* had most recent common ancestor, which was clearly shown in Fig 3 F, that consistent with Wang (Wang et al., 2013) and Li (Li Y.L. et al., 2013). However, slightly different results were obtained from trees based on 12 protein-coding mtDNA genes. The fact that the subfamily schizothoracinae was not a monophyletic group only resulted from that the *Percocypris pingi* as the sister clade of the primitive grade (100). The clade of specialized grade schizothoracine fishes were separated from the clade that included primitive grade schizothorcine fishes and *Percocypris pingi* with strong support (96), and then the highly specialized schizothoracine fishes were differentiated from the specialized grade with stronger support (100).

**Figure 2.**
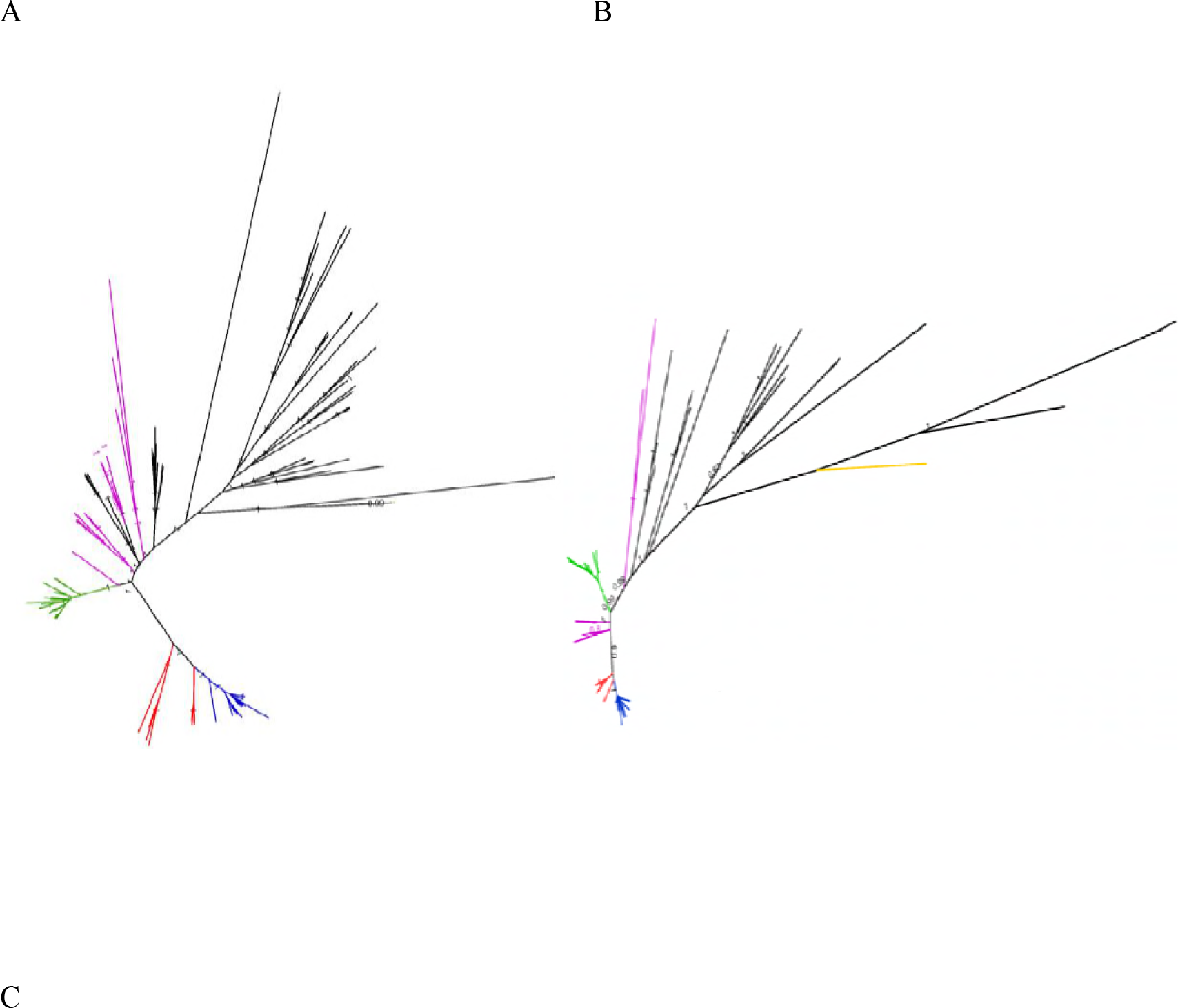
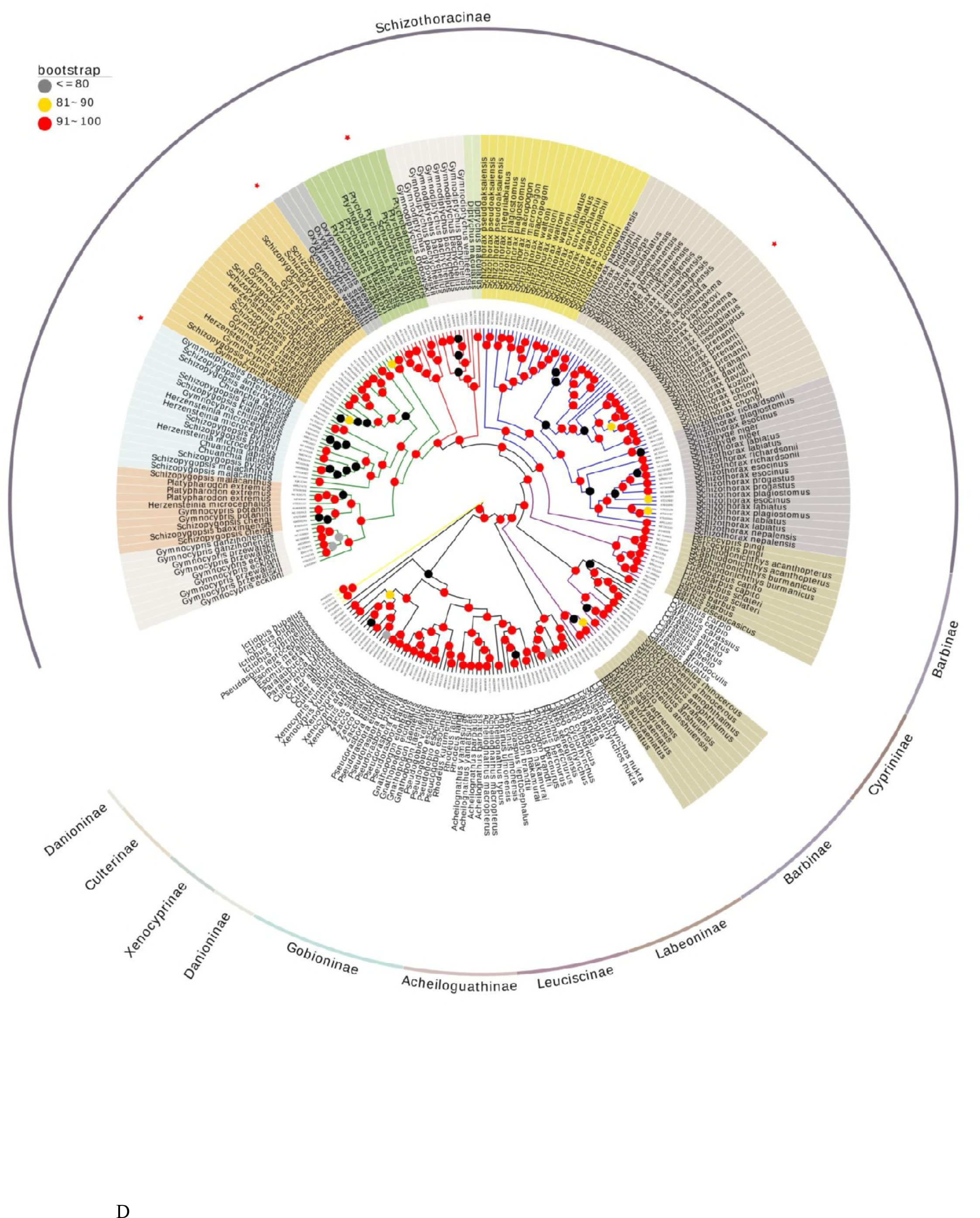
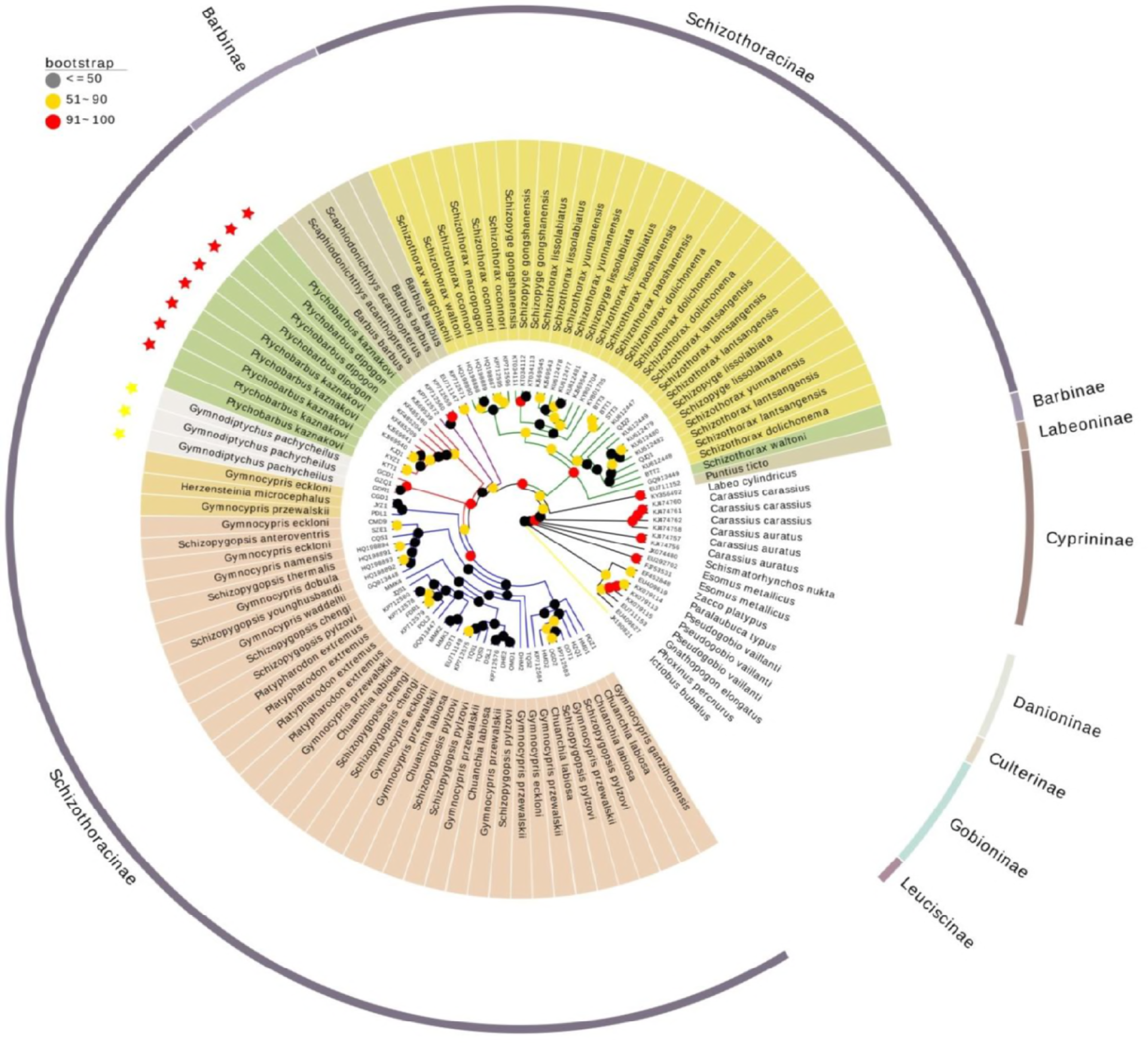
The phylogenetic tree of Cyprniformes as inferred from the mitochondrial protein-coding genes (excluding ND6 and 12SrRNA). Yellow branch represents the outgroup species, green branches represent the primitive grade of schizothoracinae, red branches represent the specialized grade of schizothoracinae and blue branches represent the highly specialized grade of schizothoracinae. A The Bayes Inferred Tree of 12 protein-coding genes based on the GTR+I+G model, Ictiobus cyprinellus was used as the outgroup. The nodal numbers indicated Bayesian posterior probability with the mcmc ngen = 2000000. B The Bayes Inferred Tree of RAG1 gene based on the GTR+I+G model, only the Ictiobus cyprinellus was used as the outgroup. The nodal numbers indicated Bayesian posterior probability with the mcmc ngen = 2000000. C The overview of Maximum Likelihood Tree based on 12 protein-coding genes with the GTRCAT model, Ictiobus were used as outgroups. The branch label numbers indicated the bootstrap probabilities (rapid bootstrap set as 1000 replications). D The overview of Maximum Likelihood Tree based on RAG1 gene with the GTRCAT model, Ictiobus were used as outgroups. The branch label numbers indicated the bootstrap probabilities (rapid bootstrap set as 1000 replications).

Both two molecular datasets were inconsistent with the conclusion of morphology, which indicated that the specialized grade originated from the primitive schizothoracinae, and the highly specialized schizothoracinae originated from the specialized schizothoracinae. Meanwhile, in phylogenetic trees based on both genes, the Schizothoracinae was divided into two major clades, indicating the subfamily schizothoracinae might have two different origins. Specialized and highly specialized Schizothoracine fishes form one clade while primitive Schizothoracine fishes made up the other major clade that indicated primitive clade was single origin and specialized and highly specialized originated from another ancestor. The primitive clade located at the bottom of the tree and the highly specialized grade lay at the top of the tree, which was in accordance with the result of He (He et al., 2004).

**Figure 3.**
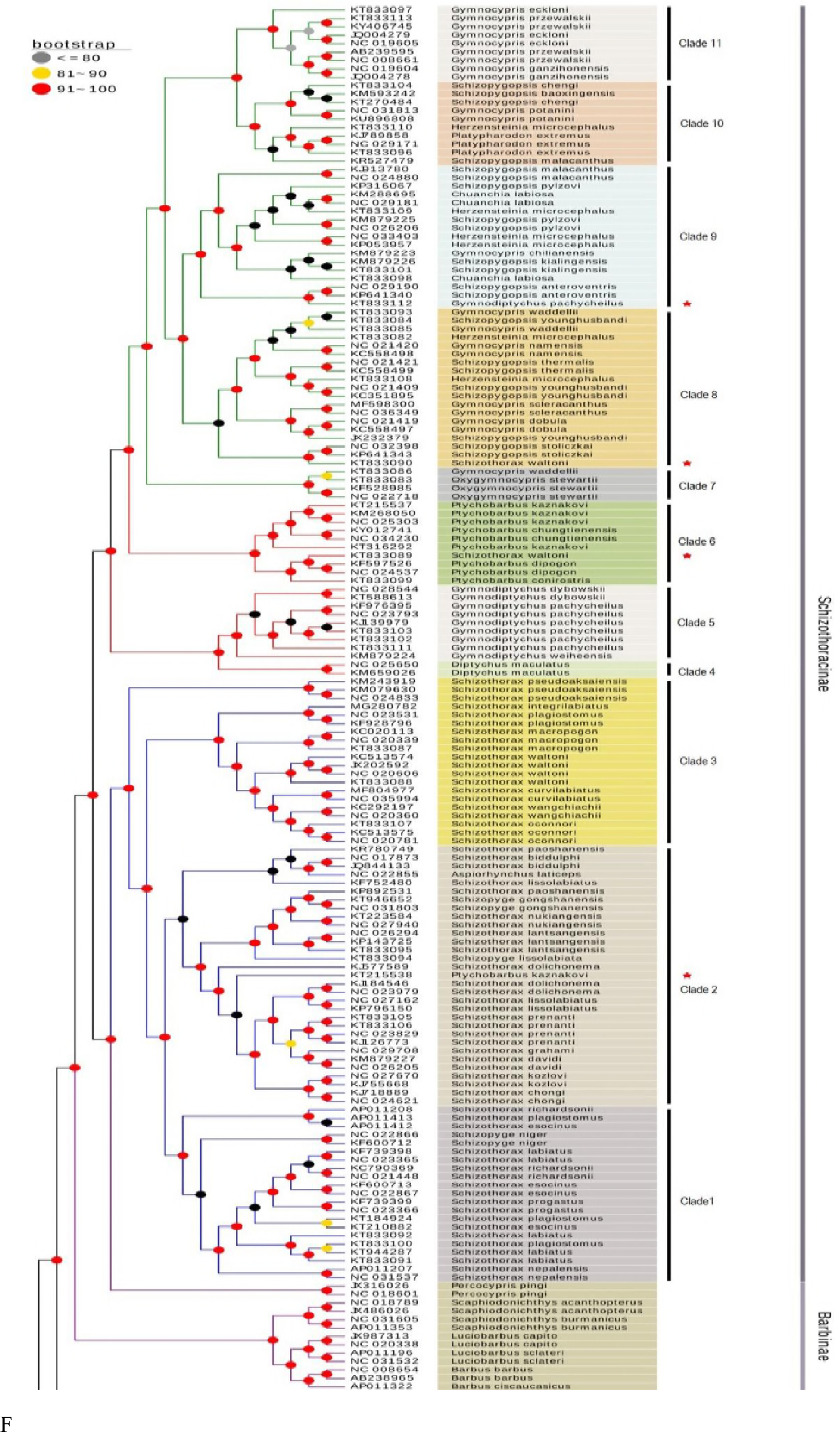
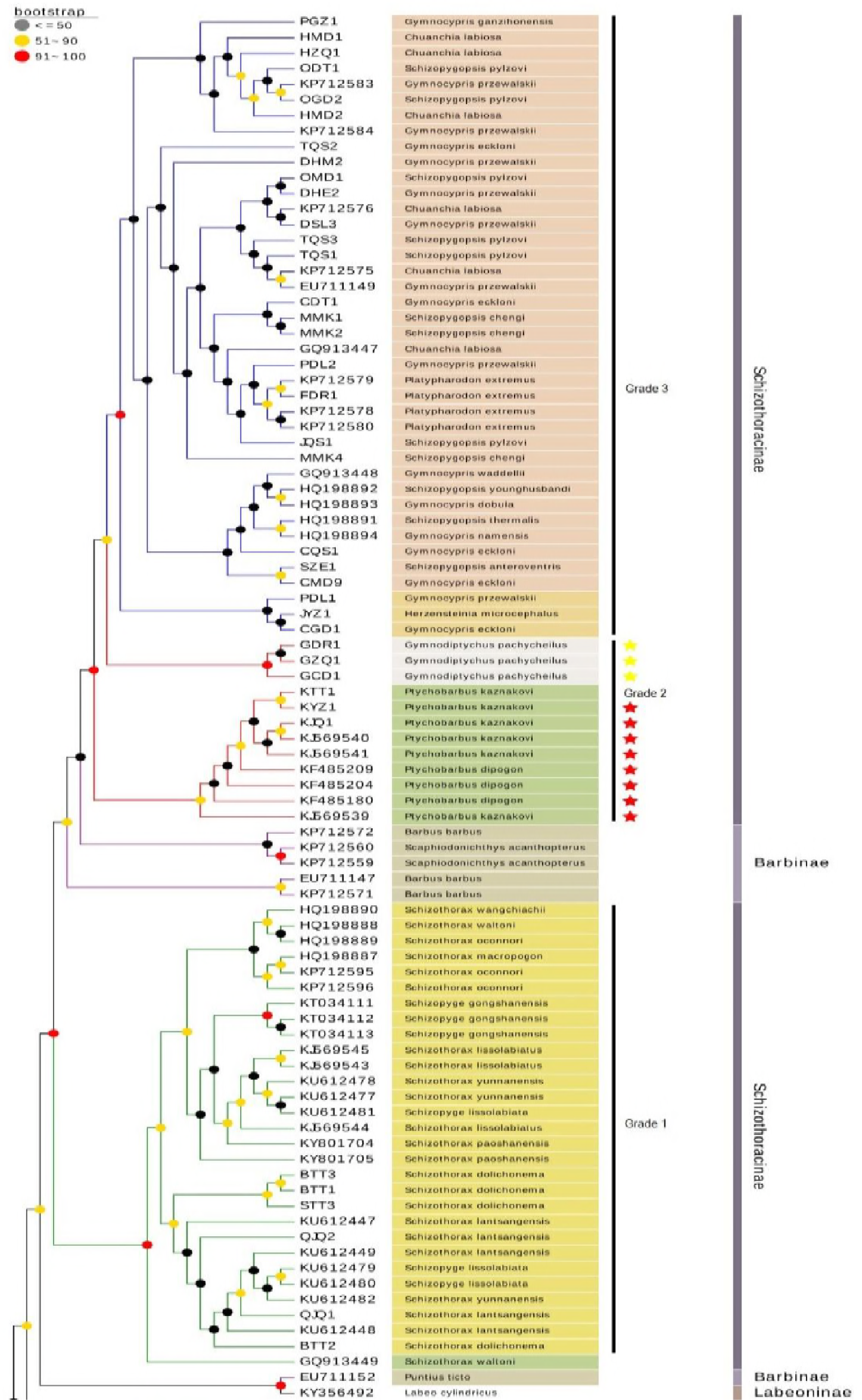
The partial enlargement of Maximum Likelihood Tree only included the subfamily schizothoracinae based on 12 protein-coding genes (E) and RAG1 gene (F). Green branches represent the primitive grade of schizothoracinae, red branches represent the specialized grade of schizothoracinae, blue braches represents the highly specialized grade of schizothoracinae, and purple branches represent the species of Barbinae. Shadows with the different color indicated different clades. Red stars in figure E represented individuals that might be incorrectly classified, while in figure F red stars and yellow stars represented genera *Ptychobarbus* and *Gymnodiptychus*.

The Schizothoracinae was comprised of 11 clades based on 12 protein-coding genes, showing in Fig 3 E, each with strong support. Clade 1 to 3 was included in primitive schizothoracinae and clade 4, 6 was the part of specialized schizothoracinae, clade 7 to 11 was involved in highly specialized schizothoracinae. It was noteworthy that the clade 6 (*Ptychobarbus*) which belonged to specialized schizothoracinae clustered with highly specialized schizothoracinae with strong support (100) that might lead us to speculate that *Ptychobarbus* was the transitional taxa that the specialized schizothoracinae evolved to highly specialized schizothoracinae and making the result be inconsistent with the monophyly of specialized schizothoracinae (Chen Z.M. & Chen, 2001), and *Diptychus* was the basal clade as monophyly. *Gymnodiptychus* clustered together as monophyly that originated from *Diptychus*. Primitive schizothoracinae was monophyletic group whereas these species of three genera tangled up with each other that strongly supposed that genus *Racoma* and genus *Schizopyge* should be merged into genus *Schozothorax*. Highly specialized schizothoracinae, Clade 7 to 11, was monophyly clustered with *Ptychobarbus* as the sister group, and *Oxygymnocypris* was the primordial genus, and phylogenetic relationships among the other genera including *Schizopygopsis, Platypharodon, Gymnocypris*, *Chuanchia*, and *Herzensteinia* were ambiguous. *Gymnocypris* possibly originated from *Oxygymnocypris. Platypharodon* clustered together as the sister group of *Gymnocypris przewalskii*. The other species of *Gymnocypris* and species of *Chuanchia* and *Herzensteinia* were embedded in genus *Schizopygopsis*. The results above were highly in line with the conclusion that members of the specialized schizothoracine group and the genera *Schizothorax*, *Schizopygopsis*, and *Gymnocypris* were paraphyletic based on complete mitochondrial genomes (Zhang J. et al., 2016). On the other hand, based on RAG1 gene (Fig 3 F), genera in highly specialized grade schizothoracinae were clustered together as a monophy clade that strongly supported (96), while genera in primitive grade and specialized grade schizothoracinae could distinguish each other clearly with unreasonable topology, which was obviously different from the results based on mtDNA genes.

### Divergence times

Our estimated divergence times of schizothoracinae based on the mitochondrial genes were much older than the estimates of He (He et al., 2004) and Ruber (Ruber et al., 2007), but was consistent with Yang (Li Y.L. et al., 2013). The divergence times of three grades Schizothoracinae were not completely consistent with three stages of the uplift of Qinghai-Tibet Plateau. The speciation time of Schizothoracinae is 66 Ma, which is consistent with the first stage of the uplift of Qinghai-Tibet Plateau. The divergence time of the primitive grade and *Percocypris pingi* was 64 Ma, while the speciation time of the primitive grade was 57 Ma. And the divergence time of specialized and highly specialized Schizothoracinae is 46 Ma, which is much older than the second stage of uplift of Qinghai-Tibet Plateau. The speciation times of specialized and highly specialized Schizothoracinae are 51 Ma and 43 Ma, respectively.

## Discussion

### Phylogeny analysis

The phylogeny of schizothoracinae is controversial all the time. Our phylogeny relationships of schizothoracinae, based on RAG1 gene, showed that the primitive grade schizothoracinae fishes clustered into a single branch that strongly supported and the specialized and highly specialized schizothoracinae clustered with some species of *Barbinae*, while phylogenetic of schizothoracinae based on 12 protein-coding mtDNA genes indicated that the specialized and highly specialized schizothoracinae as a sister clade directly clustered with clade that included the primitive grade schizothoracinae and *Percocyoris* rather than clustered with *Barbinae*. Subfamily schizothoracinae split into two clades into both molecular data, indicating that Schizothoracine fishes have two different origins, as the same as other molecular data (He et al., 2004; Li Y.L. et al., 2013; Qi et al., 2015), were inconsistent with morphological phylogenies that the schizothoracinae was monophyly and the specialized and highly specialized schizothoracinae was originated from primitive Schizothoracine fishes. In morphological phylogenies of schizothoracinaea, the trophic morphologies mainly were selected as criterions of taxonomy, such as the rows of pharyngeal teeth, lower jaws horn, pharyngeal bone, and the skull. To some extent, those morphological characters were determined by their habits and foraging ways. Convergent evolution, the same environment prompt to form the similar morphologies, enforces the different species to cluster together, making the traditional taxonomy of the schizothoracinae was different with the molecular results (Qi et al., 2012). The convergent evolution is a common phenomenon such as the lower lip of *Labeoninae* (Li Jun-bing et al., 2005), eye and pigment degeneration and well developed projection of frontal bones of the cave species (Xiao et al., 2005), the morphological similarities of ground tits and ground jays result from convergent evolution (Qu et al., 2013). Those characters of morphologies are non-homologous but are similar that are meaningless to phylogenies. And then the low level of sequence divergence make the molecular mutations don’t have enough time to stabilize and accumulate that result in the rapid differentiation of morphology don’t synchronously reflect on the molecular variations. And ancestral polymorphism, rapid evolution, expansion and diversification process, mitochondrial introgressive hybridization may also are the main factors to the inconsistence between molecular phylogenies and morphological phylogenies (He & Chen, 2007). RAG1 gene, related to immune system, has the common disadvantages of nuclear genes as molecular markers, such as heterozygous ambiguity and paralogy, few available variable sites resulting from sequences conservation (Chen W.J. et al., 2008; Li Chenghong et al., 2007; Saitoh & Chen, 2008). Therefore, the phylogenetic trees reconstructed based on the two molecular data were different, and we speculated that the phylogenetic relationships of subfamily schizothoracinae based on 12 protein-coding genes were more reliable.

Our analysis of phylogeny of schizothoracinae was also different from other some molecular phylogeny. The most prominent was that the closest relative species of schizothoracinae is *Percocypris pingi*, which was consistent with Yang (Yang et al., 2015). The genera of specialized grade are different from the other phylogeny. The *Diptychus* as the basal clade of specialized grade schizothoracinae that *Gymnodiptychus* was evolved from genus *Diptychus*, but *Ptychobarbus* was advanced genus which was clustered together with highly specialized grade schizothoracinae that were inconsistent with the result that *Gymnodiptychus* was advanced genus that evolved from *Ptychobarbus* based on Cyt *b* (Chen Z.M. & Chen, 2001). Maybe they sampled only minority species and individuals or only used Cyt *b* mitochondrial gene resulting in different conclusions. In the resolution of interrelationships amongst long-diverged groups, the length of the sequence was particularly important (Rogers, 2001). Therefore, just using the Cyt *b* gene may be inappropriate to speculate the phylogeny of schizothoracinae. Sampling as well plays a speculate role in reconstruct phylogenic relationships that we need to choose sufficient species and individuals.

### Divergence times of schizothoracinae

The divergence times of three grades of Schizothoracinae were not completely consistent with the three stages of uplift of Qinghai-Tibet Plateau, which was contradicted with the morphology results. In our data, the speciation of Schizothoracinae was 66 Ma, which was the time of the first stage of uplift of Qinghai-Tibet Plateau, but the speciation times of primitive Schizothoracinae and the specialized Schizothoracinae and highly specialized Schizothoracinae was much older than the second and third stage of uplift of Qinghai-Tibet Plateau. The speciation time of primitive clade and the specialized - highly specialized clade was very close. So the result of the divergence times was consistent with the phylogeny result, supporting that Schizothoracinae has two independent origins. The primitive clade was originated from the original genus of *Barbinae*, while the specialized and highly specialized clade, as a sister group with the clade consist of the primitive clade and *Percocypris pingi*, originated from the other genus of genus of Barbinae.

## Acknowledgements

This work was supported by the Natural Science Foundation of Qinghai Province (No. 2016-ZJ-956) and performed at Key Laboratory of Medicinal Plant and Animal Resources of Qinghai-Tibetan Plateau in Qinghai Province (2017-ZJ-Y13), Key Laboratory of Evolution and Adaptation of Plateau Biota, and Key Laboratory of Animal Genomics in Qinghai Province (Northwest Institute of Plateau Biology, Chinese Academy of Sciences, 2017-ZJ-Y23).

## Author contributions

All authors collected samples together. Yurong Du, Jianbin Ma and Songchang Guo planned the experiments and helped with analyzing the paper; Delin Qi helped with species classification. Ting Wang analyzed the data and wrote the paper. Ting Wang and Juan Chen performed the experiments, Shihao Sun and Zhonghao Liu assisted with the experiments. Songchang Guo edited the paper. The authors alone are responsible for the content and writing of this paper. The authors declare no conflicts of interest.

